# Evidence of a noncoding transcript of the *RIPK2* gene overexpressed in head and neck tumor

**DOI:** 10.1101/466011

**Authors:** Ulises M. M. Villagra, Bianca R. da Cunha, Giovana M. Polachini, Tiago Henrique, Carlos H. T. P. da Silva, Olavo A. Feitosa, Erica E. Fukuyama, Rossana V. M. López, Emmanuel Dias-Neto, Fabio D. Nunes, Patricia Severino, Eloiza H. Tajara

## Abstract

Receptor-interacting proteins are a family of serine/threonine kinases, which integrate extra and intracellular stress signals caused by different factors, including infections, inflammation and DNA damage. Receptor-interacting serine/threonine-protein kinase 2 (RIP-2) is a member of this family and an important component of the nuclear factor NF-kappa-B signaling pathway. The corresponding human gene *RIPK2* generates two transcripts by alternative splicing, the full-length and a short transcript. The short transcript has a truncated 5’ sequence, which results in a predicted isoform with a partial kinase domain but able to transduce signals through its caspase recruitment domain. In this study, the expression of *RIPK2* was investigated in human tissue samples and, in order to determine if both transcripts are similarly regulated at the transcriptional level, cancer cell lines were submitted to temperature and acid stresses. We observed that both transcripts are expressed in all tissues analyzed, with higher expression of the short one in tumor samples, and they are differentially regulated following temperature stress. Despite transcription, no corresponding protein for the short transcript was detected in tissues and cell lines analyzed. We propose that the shorter transcript is a noncoding RNA and that its presence in the cell may play regulatory roles and affect inflammation and other biological processes related to the kinase activity of RIP-2.

## Introduction

Unicellular and multicellular organisms are constantly exposed to stressful environments. Chemical and physical stimuli trigger different adaptive responses, which will determine the capability of the organism to maintain internal homeostasis [1].

Receptor-interacting proteins (RIP) are a family of serine/threonine kinases, which integrate extra- and intracellular stress signals and share a homologous kinase domain at the N-terminus, but have different C-terminal functional domains [2-4]; RIP-2 (receptor-interacting serine/threonine-protein kinase 2) is a member of the RIP family, which has received attention in the recent years for its role in modulating immune and inflammatory processes [5], and as a sensor of cellular stress [6]. It is expressed at high levels in several normal human tissues [7], as well as in pathological conditions, for example ulcerative colitis [8], triple-negative breast cancers [9,10] and in stressful conditions, such as after hypoxic/ischemic insults [11]. Conversely, lower levels of RIP-2 have been correlated with tumor progression in squamous cell carcinoma (SCC) of the oral cavity [12].

RIP-2 is the only member of the RIP family that besides phosphorylating serines and threonines is able to autophosphorylate tyrosine residues [13,14]. Its ATP- and substrate binding sites spread over much of the N-terminal kinase domain, and a caspase recruitment domain (CARD) is present in the C-terminal region [15] (isoform 1, Fig 1A). CARD specifically interacts with the nucleotide-binding oligomerization domain-containing protein 1 (NOD1) and NOD2 (also called CARD-4 and CARD-5, respectively), which are intracellular receptors for innate immunity and involved in sensing the presence of pathogens. After activation by bacterial peptidoglycans, NOD1 and NOD2 associate with RIP-2 via CARD-CARD interaction and promote the expression of immune response and inflammatory genes through the nuclear factor-kappa B (NFKB) signaling [16]. NOD1 and NOD2 also cooperate and share redundant roles with Toll-like receptors (TLRs) in detecting bacteria, but there’s no consensus on the participation of RIP-2 in TLR signaling (reviewed by [3]). Recently, it has been demonstrated that RIPK2 kinase activity and auto-phosphorylation are not required for NOD2 inflammatory signaling. In fact, NOD2 pathway activation and cytokine production depends on RIP-2 polyubiquitination at several lysines, a process relied on the ubiquitin ligases:RIP-2 kinase domain interaction. Thus, although the kinase domain is not functionally important for NOD2 signaling, it is correlated with NOD2 activation since RIP-2 auto-phosphorylation creates a substrate for ubiquitin ligase binding [17].

**Fig 1.**
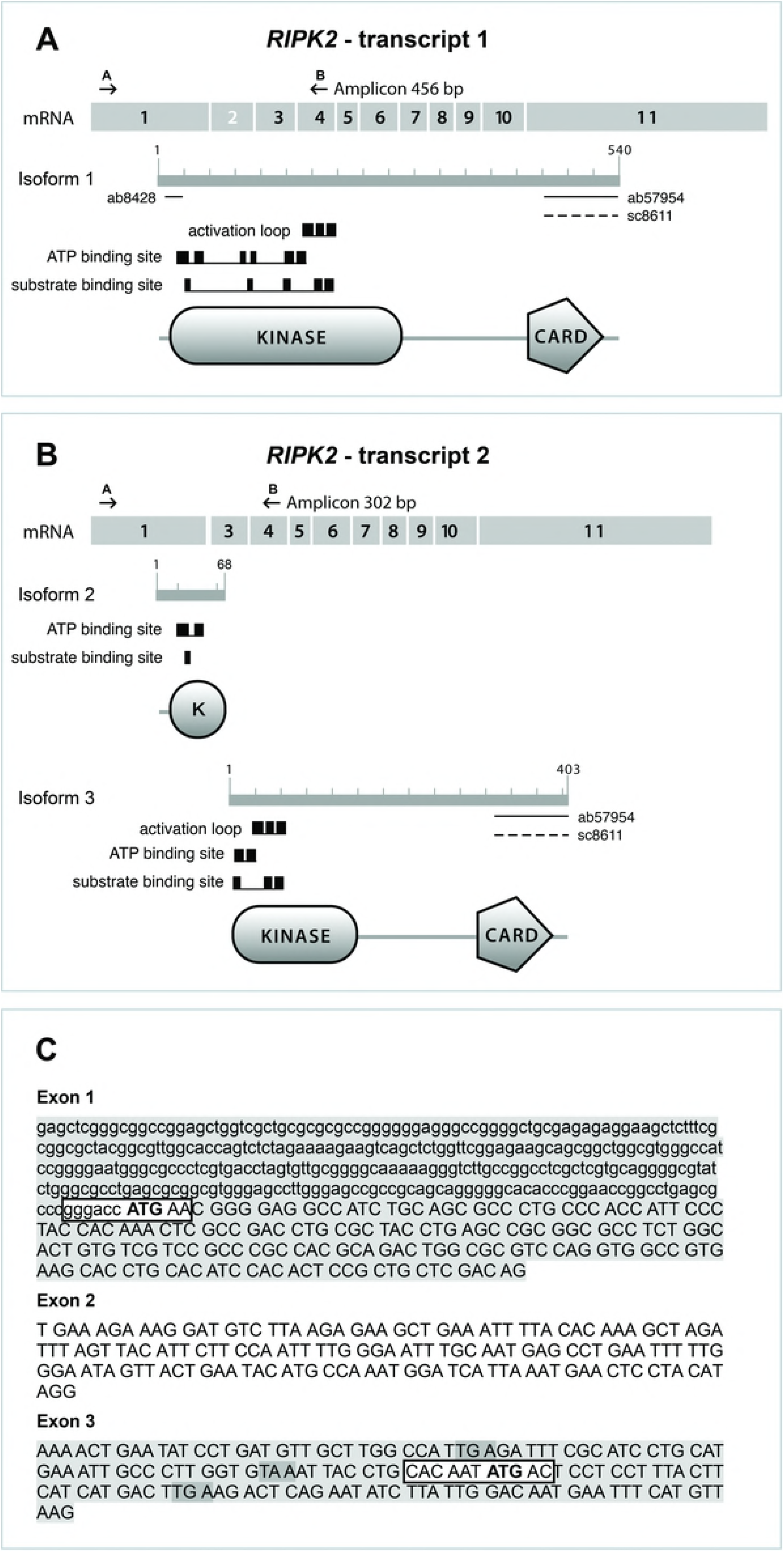
Diagrams of RIPK2 splice transcripts. (**A**) The transcript 1 encodes the longer isoform and (**B**) the transcript 2 presents skipping of exon 2 and encodes a very short isoform 2 [18] and a predicted isoform 3. Arrowheads indicate the positions of the forward primer A and reverse primer B for RT-PCR expression analysis, and horizontal bars below the isoform 1 and 2 indicate the epitope region for anti-RIP-2 ab8428, ab57954 and sc8611 used in the present study. (**C**) 5’UTR (lower case) and codons (capital letters) of exons 1, 3 (gray) and 2 (white). Kozak sequences in boxes. First ATGs of full-length isoform and predicted isoform 3 in bold. Premature stop codons generated by the frameshift due to exon 2 skipping=dark gray. Kinase/K=protein kinase domain; CARD=caspase recruitment domain; activation loop; ATP binding site; substrate binding site; according to Batch Conserved Domain-Search at NCBI.

The full-length human *RIPK2* transcript (GenBank accession number NM_003821.5), here named transcript 1 (Fig 1A), has 2588 bps and is composed of 12 exons spanning 33-kb of genomic sequence on chromosome 8q21. In 2004, we suggested an alternative splicing for *RIPK2* transcribing a short variant (AY562996, currently included within the predicted transcript XM_005251092.3) [19], as depicted in Fig 1B. This variant, here named transcript 2, has 2389 bps and derives from the skipping of exon 2 (154 bps), which alters the reading frame producing several premature stop codons. However, a potential translation initiation codon AUG (nucleotides 85-87 of exon 3) is in-frame with the downstream *RIPK2* sequence, hence with no subsequent premature termination codon (Fig 1C). Translation from this codon may give rise to an amino-terminal truncated protein (XP_005251149.1, isoform 3 in this study, with 403 residues, predicted molecular weight of 45,582 Da) lacking the first 137 amino acids of RIP-2 isoform 1 (NP_003812.1).

Alternatively spliced transcript 2 was also studied by Krieg and collaborators [18], who reported a protein product (isoform 2, Fig 1B, top) with extensive truncation of the N-terminal kinase domain and a complete lack of the intermediate domain and CARD due to a frame shift generating a premature stop codon. Krieg et al. also reported that this isoform of RIP-2 lacks the biological effects described for the isoform 1. We here investigated if the use of the downstream alternative translation initiation site may generate an isoform 3 that would keep the original C-terminal structure including CARD, but would present a truncated kinase domain (Fig 1B, bottom), with potential consequences for protein function, and cellular localization if the localization signals were also deleted.

Since RIP-2 kinase integrates extra and intracellular stress signals and modulates immune responses [5], we reasoned that physiological and environmental changes, such as hyperthermia and acid stress caused by infections and inflammatory processes, might affect the expression of their transcripts and, therefore, could lead to changes in levels of the isoforms depicted in Fig 1. The use of alternative splicing sites may differ among cell types and phases of development [20-23], or be associated with stress conditions, such as temperature stress [24] and oxidative stress [25].

In the present study, the expression of *RIPK2* transcripts and protein products was evaluated in normal human tissues and in SSC samples and, in order to investigate if they are regulated in response to stress conditions, we analyzed their expression upon heat/cold and acid stress in human cancer-derived cell lines.

## Material and methods

### Samples and cell lines

Nine samples of normal human tissues removed at autopsy (brain, testis, heart, lung, stomach, kidney, larynx, liver and tongue) and 16 matched tumor/resection margin samples of oral SCC were used to evaluate the expression of the two transcripts of the *RIPK2* gene. RIP-2 protein levels were analyzed in another set of 19 matched tumor/resection margin of oral and laryngeal SCC.

For the stress experiments, we used the human cell lines FaDu (HTB-43, derived from SCC of the hypopharynx), and SiHa (HTB-35, derived from cervix SCC). The cell lines were cultured in Minimum Essential Medium (MEM, 552, Cultilab), supplemented with 10% fetal bovine serum (FBS, 63, Cultilab), 10 mM non-essential amino acids (M7145, Sigma), 2 mM L-glutamine (687, Cultilab), 1 mM sodium pyruvate (P5280, Sigma), 1.5 g/L sodium bicarbonate (S5761, Sigma), penicillin (100 units/mL) and streptomycin (90 μg/mL) (1012, Cultilab), in a humidified atmosphere with 5% CO_2_ at 37°C. The study protocol was approved by the National Committee of Ethics in Research (CONEP 1763/05, 18/05/2005, and CONEP 128/12, 02/03/2012) and informed consent was obtained from all patients enrolled.

### Temperature and acid stresses

Prior to stress experiments, cell lines were grown to 80-90% confluence and cell cycle synchronized in serum-free medium for 24 h. For temperature stress, cells were maintained in medium plus 10% FBS at 40°C, 17°C or 5°C for 3 h (eight replicates for each condition). Six control replicas were also cultured in medium plus 10% FBS at 37°C for 3 h. Acidic shock was performed by maintaining the cultures (four replicas) in an atmosphere with elevated CO_2_(10% CO_2_) for 24 h or 72 h. Four control replicas were cultured in a humidified atmosphere with 5% CO_2_ at 37°C for the same time period. After the incubation period, cells were immediately lysed by adding TRIzol (15596026, ThermoFisher), and stored at −80°C until RNA extraction.

### RNA extraction and cDNA synthesis

Total RNA from tissue samples and cell lines was obtained following the TRIzol protocol. Integrity of the RNA was confirmed by agarose gel electrophoresis, and the purity and concentration were determined using the NanoDrop ND-1000 spectrophotometer (Thermo Fisher). One microgram of total RNA was converted to cDNA using the High Capacity cDNA Reverse Transcription kit (4368813, Thermo Fisher), according to the manufacturer’s instructions.

### Detection of RIPK2 transcripts by polymerase chain reaction

The PCR amplification of *RIPK2* transcripts was performed using the oligonucleotides 5’-CGCCTCTGGCACTGTGTCGT-3’ (forward primer A) and 5’-CGTGACTGTGAGAGGGACAT-3’ (reverse primer B). The PCR reaction was carried out in a total volume of 25 μL containing 1X PCR buffer, 1 mM MgCl_2_, 2 μM of each *RIPK2* primer, 2 μM *GAPDH* primers, 5 mM dNTPs mix, 1 U Taq DNA polymerase (EP0402, ThermoFisher) and 50 ng of cDNA. After pre-incubation for 5 min at 94°C (initial denaturation), the amplification was carried out through 35 cycles at 94°C for 50 s, 58°C for 40 s, 72°C for 50 s, and 72°C for 10 min, using a thermal cycler (9700 GeneAmp PCR System, Applied Biosystems). PCR primers for the endogenous control gene *GAPDH* were GAPDHF (5’-ACCCACTCCTCCACCTTTGA-3’) and GAPDHR (5’-CTGTTGCTGTAGCCAAATTCGT-3’). The expected lengths for PCR amplicons were 101 base pairs (bps) for *GAPDH* and 456 or 302 bps for *RIPK2* transcripts. Amplicons were separated on 2% agarose gels, bands were quantified by densitometry using Image J software, and sequenced in both directions after being isolated from the gels. The sequences were analyzed using BLAST similarity search against the non-redundant database available from the National Center for Biotechnology Information (NCBI) [26].

### Evaluation of RIPK2 transcripts by relative quantification using RT-qPCR

The expression of *RIPK2* transcripts in matched tumor/resection margin samples and in cell lines following stress treatment was investigated by quantitative PCR (qPCR). Reactions were performed in triplicate using an ABI Prism 7500 Sequence Detection System (Applied Biosystems). The primers were manually designed and optimized for RT-qPCR using basic parameters for PCR primer design. The final sequences were 19-24-bp long, with 30-70% GC content and producing a short amplicon size (66-104 bps), as follows: *RIPK2* transcript 1 forward 5’-AGAAGCTGAAATTTTACACAAAGC-3’ and reverse 5’-CCATTTGGCATGTATTCAGTAAC-3’; *RIPK2* transcript 2 forward 5’-TGCTCGACAGAAAACTGAATATC-3’ and reverse 5’-AAGGAGGAGTCATATTGTGCAG-3’; *GAPDH* forward 5’-ACCCACTCCTCCACCTTTGA-3’ and reverse 5’-CTGTTGCTGTAGCCAAATTCGT-3’; *TUBA1C* forward 5’-TCAACACCTTCTTCAGTGAAACG-3’ and reverse 5’-AGTGCCAGTGCGAACTTCATC-3’; *ACTB* forward 5’-GGCACCCAGCACAATGAAG-3’ and reverse 5’-CCGATCCACACGGAGTACTTG-3’. All primers were purchased from Invitrogen. Briefly, reactions were carried out in a total volume of 20 μL, with 10 μL SYBR Green PCR Master Mix (4385612, ThermoFisher), 250 nM of each primer and 20 ng cDNA. The PCR conditions were 50°C for 2 min, 95°C for 10 min, followed by 40 cycles of 95°C for 15 s, 58°C for 10 s, 60°C for 1 min. Following PCR, dissociation curve analyses were performed to confirm the single gene product. Adequate internal control reference genes were selected using the geNorm algorithm [27] and *TUBA1C* (stress assays) and *ACTB* (tumor/margin samples) were selected. The relative expression ratio (fold-change) of the target genes was calculated according to Pfaffl [28]. Statistical analysis was carried out by a two-tailed unpaired t test using GraphPad Prism (GraphPad Software). Values were Log2 transformed and those below −1 indicated down-regulation in gene expression while values above 1 represented up-regulation in test samples compared with control samples.

### Protein sequence alignment and homology modeling procedures

Homology modeling of RIP-2 isoforms 1 and 3 was carried out using the MODELLER software that performs modeling by satisfaction of spatial restraints [29]. Six homologues with structures available in the Protein Data Bank (codes 2GSF, 1JPA, 1K2P, 1U59, 1UWH, 2EVA) used as templates were selected through a non-redundant BLASTp search [26]. Two putative conserved domains with statistical significance were detected: TyrKc and S_TKc, which correspond, respectively, to the catalytic domains of tyrosine and serine/threonine protein kinases, and include the leucine L10-threonine T296 sequence. These six templates share sequence identities of 27.1% (Eph receptor tyrosine kinase, PDB code 1JPA) to 30% [Transforming growth factor-beta (TGF-beta)-activated kinase 1 - TAK1, PDB code 2EVA] with RIP-2. Analyses were performed using pairwise alignments via the AMPS (Alignment of Multiple Protein Sequences) package [30]. Previous to the modeling, a final multiple alignment was obtained by analyzing the superposition of the six structures regarding the alpha-carbons of the residues, using the INSIGHT II program, version 2005 (Accelrys Inc, San Diego, CA, USA), which allowed refine the previous alignment obtained from the AMPS MULTALIGN module. The information of secondary structure in the template sequence was incorporated into this previous alignment using the MULTALIGN module of the AMPS package, with the restriction that all insertions and deletions were limited to regions outside the common core of alpha-helices and beta-sheets. A gap penalty of 1000 was fixed to any deletion or insertion inside a secondary structure element. The alignment obtained was edited, investigating and considering the aligned residues, which were close in the space, as visualized in the structural superposition. This procedure resulted in a final alignment that is different from the one based on the Dayhoff matrix (PAM 250) used in AMPS.

NetPhosK 1.0 server [31] was used for phosphorylation site analyses. The algorithm produces neural network predictions of kinase-specific eukaryotic protein phosphorylation sites. Currently, NetPhosK covers the following kinases: PKA, PKC, PKG, CKII, Cdc2, CaM-II, ATM, DNA PK, Cdk5, p38 MAPK, GSK3, CKI, PKB, RSK, INSR, EGFR, and Src.

### Western blot

Western blot analysis aimed at detecting isoforms 1 and 3. The antibodies used were: (a) polyclonal anti-RIP2 (ab8428, Abcam), immunogenic peptide corresponding to amino acids 11/30 of human RIP-2 (which are only present in isoform 1), N-terminal domain, diluted 2:1000 or 3:1000; (b) monoclonal anti-RIP2 (ab57954; Abcam), immunogenic peptide corresponding to amino acids 431-541, C-terminal domain, 3 μg/mL; (c) polyclonal anti-RICK (C-19) (sc8611, Santa Cruz), immunogenic peptide corresponding to C-terminal domain according to the manufacturer’s datasheet, diluted 1:200; (d) monoclonal anti-beta-actin (A5441, Sigma-Aldrich) diluted 1:5000. The antibodies mapping at C- and N-terminus of RIP-2 are depicted in Fig 1 (isoforms 1 and 3).

In brief, protein samples (30 μg) were loaded onto 12% resolving gel with 5% stacking gel (SDS-PAGE) in denaturing conditions at 120V for 80 min. The molecular weight ladder used was the PageRuler Prestained Protein Ladder (#26616; Thermo Scientific). The proteins were then transferred electrophoretically (162.5 mA per blot 70 min; Mini Protean 3 Cell, BioRad) to polyvinylidene fluoride (PVDF) membrane (IPVH00010, Immobilon-P, Millipore) with transfer buffer (25 mM Tris, 0.2 M glycine, 20% v/v methanol; Merck, Germany). Western blotting was performed using the Amersham ECL Select Western Blotting Detection Reagent (RPN2235, GE Healthcare, Life Sciences), according to the manufacturer’s protocol. The immunoreactive proteins were visualized using horseradish peroxidase-coupled secondary antibody (074-1506, 074-1806, KPL, Kirkegaard & Perry Laboratories Inc., Gaithsburg, MD, USA) and enhanced chemiluminescence reagent (Amersham ECL Select kit, RPN2235, GE Healthcare). The Fusion FX5 system (Vilber Lourmat) was used for the acquisition of the signal. The PVDF membranes were also submitted to chromogenic staining using the Western Breeze kit (Invitrogen). The blots were then scanned and analyzed (Gel Logic HP 2200 imaging system; Carestream Health Inc./Kodak Health Group, Rochester, NY, USA).

## Results

### Identification and expression patterns of *RIPK2* transcripts

Alternatively spliced transcripts 1 and 2 of *RIPK2* were co-expressed in normal tissue samples from brain, testis, heart, lung, stomach, kidney, larynx, liver and tongue (Fig 2A). Both transcripts were also detected in tumor and tumor free surgical resection samples from patients with oral SCC as illustrated by 5 matched tumor/surgical margin samples in Fig 2B. Quantitative real time PCR showed no difference in expression between full length transcript 1 and their respective surgical margins, whereas a significant higher level of shorter transcript 2 was observed (p=0.03) (as illustrated by 11 paired samples in Fig 2C).

**Fig 2.**
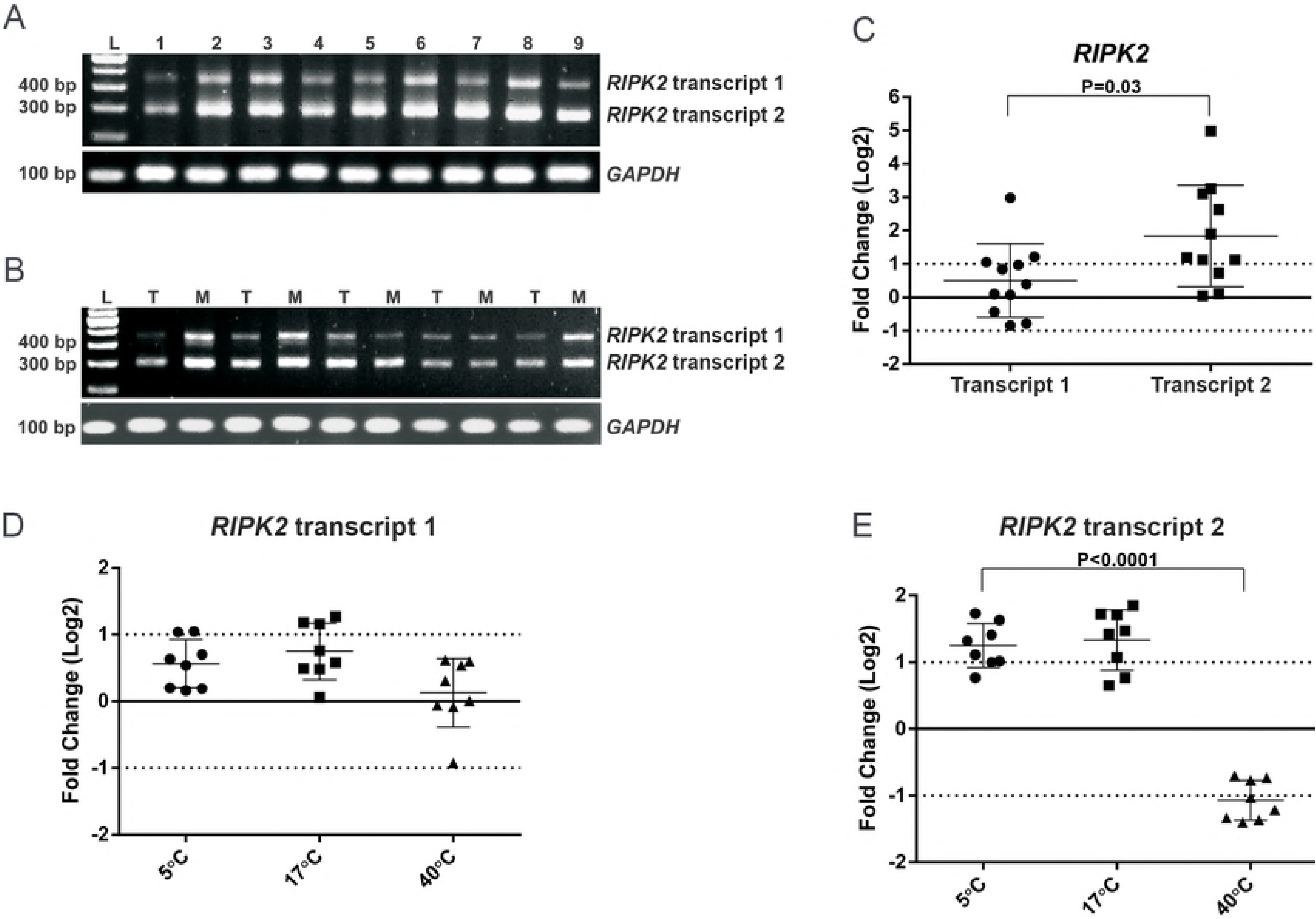
*RIPK2* mRNA expression in normal tissues, oral SCC samples and cell lines under stress conditions. (**A-B**) Conventional PCR products from *RIPK2* transcript 1 (456 bps), transcript 2 (302 bps), and *GAPDH* (101 bps) in: (**A**) normal human tissues: 1=brain, 2=testis, 3=heart, 4=lung, 5=stomach, 6=kidney, 7=larynx, 8=liver, 9=tongue; (**B**) samples from patients with oral cancer: T=tumor; M=resection margin; L=100-bp fragment size marker. (**C-E**) RT-qPCR products. (**C**) Log2 fold-change of *RIPK2* transcripts showing that transcript 2 has a higher expression than transcript 1 in tumors normalized with matched resection margins (p=0.03, unpaired t test). *ACTB* was used as the expression reference. (**D**) Expression of *RIPK2* transcript 1 and (**E**) of transcript 2 in FaDu cells maintained at 5°C, 17°C or 40°C for 3 h, normalized with control cells at 37°C (calibrator sample). Temperature stress induced a significant increase in transcript 2 expression level at lower temperatures and a decrease at a higher temperature (p<0.0001, unpaired t test), but no effect on transcript 1 expression. *TUBA1C* was used as the expression reference. Values were log2 transformed (y-axis) so that all values below −1 indicate down-regulation in gene expression while values above 1 represent up-regulation. The error bar represents the mean ± S.E.M (standard error of the mean). Significant differences: p<0.05.

### Splicing patterns of *RIPK2* under stress conditions

To investigate whether alternative splicing of *RIPK2* is induced or inhibited by stress conditions, two cell lines (FaDu and SiHa) were exposed to severe temperature stress (40°C, 17°C and 5°C), and acid stress (atmosphere of 10% CO_2_). Acid stress resulted in no effect on the expression of both transcripts (data not shown), whereas heat/cold stress induced a significant increase in shorter transcript 2 expression level at lower temperatures and a decrease at a higher temperature (p<0.0001), but no effect on full-length transcript 1 expression. This result was only observed in FaDu cells and suggests that distinct regulatory mechanisms may interfere in the alternative splicing of *RIPK2* and that this may be tissue or context dependent.

### Protein sequence alignment and homology modeling procedures

The model built for the catalytic domain of isoforms 1 and 3 showed good stereochemical quality (Fig 3 – kinase domain present only in isoform 1 inside dashed-box; superposition of the isoform 1 and 3 models outside the box). Despite overall low sequence identity among the complex structures of the homologue RIP-2 proteins, the active sites are structurally similar and reasonably well conserved. The model built for the isoform 1 is 137 residues larger than the one obtained for the isoform 3, and includes leucine L10 to threonine T296 of the overall sequence. Regarding the phosphorylation sites predicted for the isoform 1, the NetPhosK 1.0 server pointed out 10 sites, which are absent in the isoform 3 sequence: serines at positions 8, 25, 29, 33, 58, 76, 102, and threonines at positions 12, 31, 95. The study of Dorsch’s group [32] suggested that serine S176 is an important autophosphorylation site for RIP-2, and this phosphorylation can be used to monitor the activation state of RIP-2. Fig 3 shows the superposition of the two models, as well as the localization of serine S176, which seems to be conveniently accessible to the solvent and to phosphorylation. The lysine K47 in the conserved ATP-binding site and the critical polyubiquitination site lysine K209 are also shown in Fig 3.

**Fig 3.**
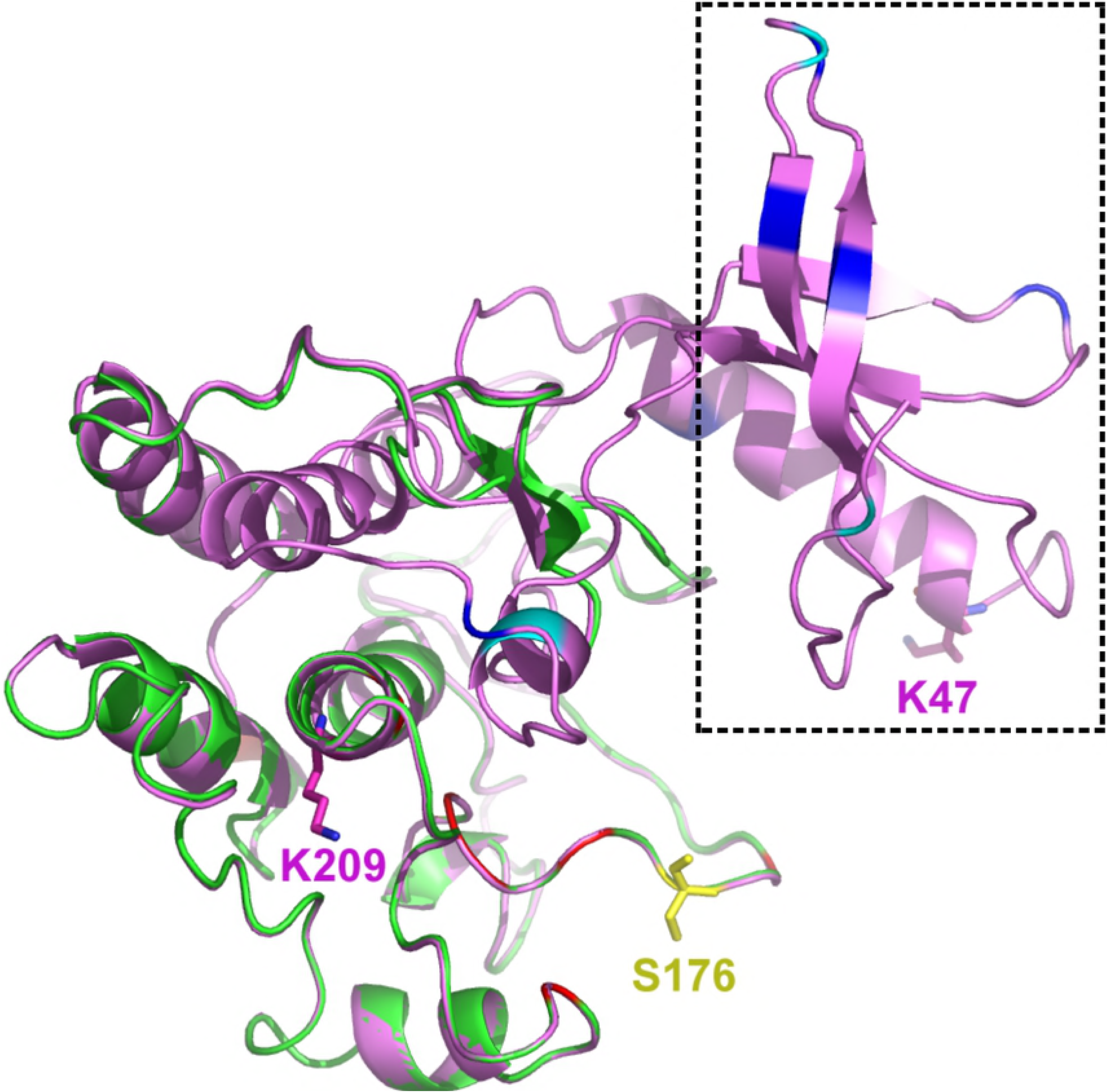
Homology modeling of the catalytic domain of RIP-2 isoforms. Superposition of the models built for the RIP-2 isoforms 1 and 3 (ribbon diagram colored in magenta and green, respectively), with the kinase domain highlighted by the dashed-box (present only in isoform 1), and the superposition of the isoform 1 and 3 models outside the box. Isoform 3 lacks the first 137 amino acids of RIP-2 and, consequently, the residue critical for kinase activity of RIP-2 (lysine K47). Ten phosphorylation sites (serine and threonine residues, respectively) predicted for isoform 1 and absent in isoform 3 sequence are indicated in blue and cyan inside the dashed box. Serine S176 is shown in yellow stick, and is indicated to be conveniently accessible to the solvent and to phosphorylation. Lysines K47 and K209 are also shown in colored sticks.

### Immunodetection of RIP-2 isoforms

Western blot analysis detected RIP-2 protein in extracts derived from FaDu cell line and from human tumors and their resection margins (Fig 4). The best results were obtained with the N-terminal domain anti-RIP2 ab8428 and C-terminal anti-RICK sc8611 antibodies, whereas ab57954 antibody yielded weak or non-specific bands. In cell line samples, Western blot demonstrated a single immunoreactive band at ~61 kDa, consistent with the molecular weight of the isoform 1 (61,195 Da). When cells were exposed to heat/cold stress, a lower expression of the RIP-2 isoform 1 was observed in FaDu cells maintained at low temperatures compared with control cells at 37°C (Fig 4A).

**Fig 4.**
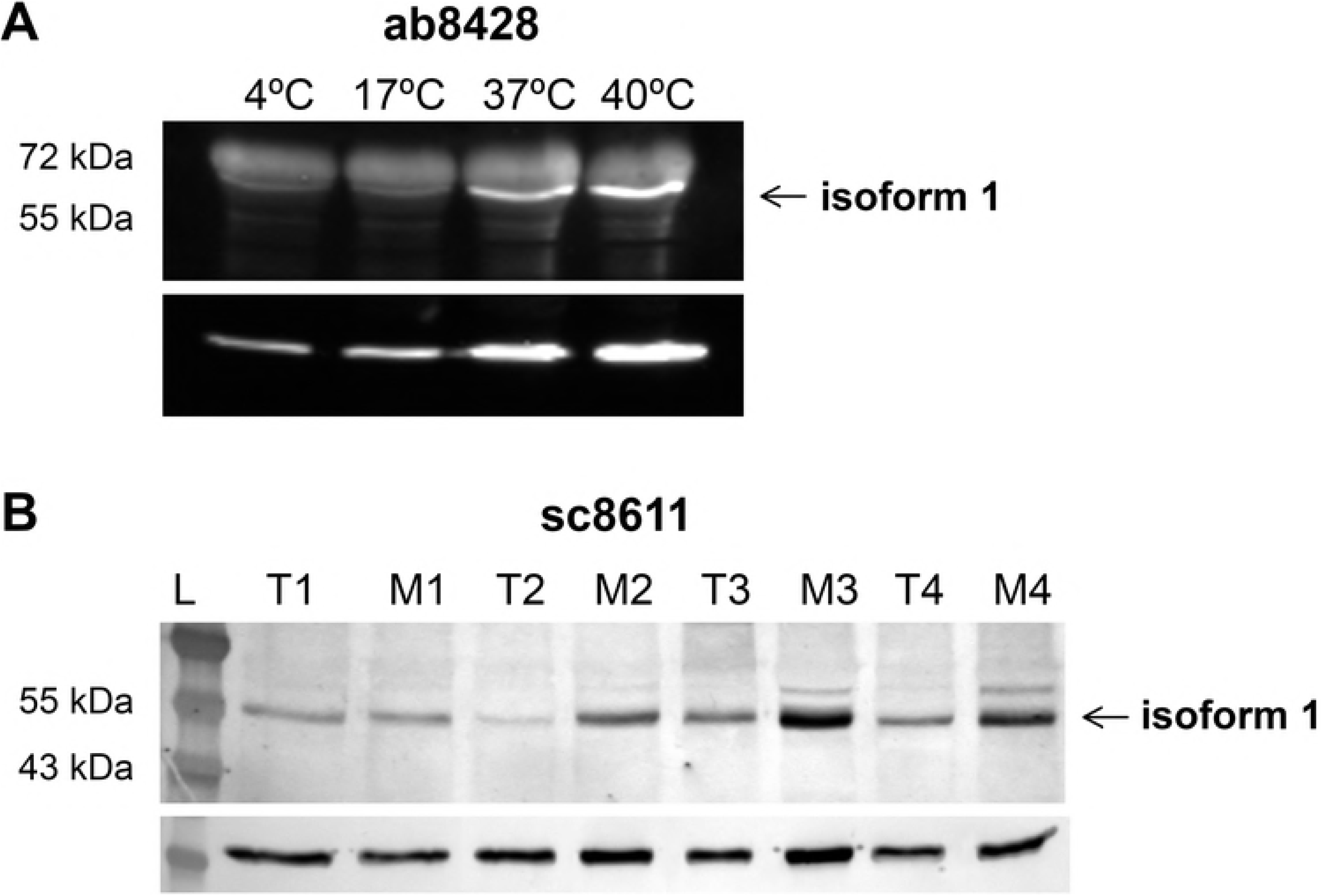
RIP-2 expression in cell lines under stress conditions and in oral SCC samples. Western blot illustrating (**A**) lower expression of the RIP-2 isoform 1 (~61 kDa) in FaDu cells maintained at low temperatures compared with control cells at 37°C (anti-RIP-2 ab8428 against N-terminus); and (**B**) an apparent decreased expression of the RIP-2 isoform 1 in tumor (T) than in resection margin (M) samples (anti-RIP-2 sc8611 against C-terminus). (**A-B**) No band that corresponds to isoform 3 was observed. Data were normalized by beta-actin. L = Protein Ladder.

Isoform 3 (predicted molecular weight of 45,582 Da) was not detected both in cell lines and normal or neoplastic tissue samples (Fig 4A, 4B), even after mass spectrometry analysis of Western blot band corresponding to the region around 45 kDa (data not shown). The result indicates that, despite the detection of transcripts 1 and 2 and the proteins predicted *in silico* for the isoforms, alternatively spliced transcript 2 does not seem to be translated into protein in normal human tissues, cancer samples or cell lines analyzed.

## Discussion

In the present study, *RIPK2* transcripts 1 and 2 were co-expressed in normal tissue samples from brain, testis, heart, lung, stomach, kidney, larynx, liver and tongue. Our stress experiments showed that one of the transcripts (transcript 2) is up-regulated at low temperatures compared with the control group (at 37°C), whereas the opposite occurs at 40°C. We tested two distinct SCC-derived cell lines, but this effect was seen only in FaDu cell line. This finding may be tissue/context dependent. In fact, literature has already referred that a mild hypothermic condition appears to induce or to inhibit synthesis of specific proteins when compared with control cells, an effect that is cell line dependent (reviewed by [33]. It’s well known that both prokaryotic and eukaryotic organisms respond to cold stress reducing transcription, translation, and metabolic processes, except in the case of cold-shock proteins [34].

Many studies are available on alternative splicing regulation by temperature and other extrinsic agents. For example, Gemignani and collaborators [35] also observed a shift in splicing of a mutated human b-globin gene affected by temperature *in vitro*, which led the authors to propose temperature changes as a treatment for β-thalassemia. More recently, Farashahi Yazd’s group [24] described a novel spliced variant of *OCT4* gene significantly elevated under heat-stress conditions, and proposed a potential role of OCT4B1 transcript and protein in mediating temperature response.

Yan and collaborators [36] found evidence that hyperthermia induces Toll-like receptors expression and TLR signaling-mediated activation of NFKB and MAPK pathways, resulting in increased synthesis of pro- and anti-inflammatory cytokines. These data suggest that fever may modulate innate immune responses by TLR pathway and, although the role of *RIPK2* in this signaling remains controversial [37-40], they provide a possible link to *RIPK2* expression changes depending on the temperature variation.

Considering these data, we hypothesize that, under stress conditions, a putative mechanism may induce higher or lower expression of the exon 2-containing transcript of *RIPK2*. Since the presumed isoform 3 has a truncated kinase domain but is potentially able to mediate NFKB activation via CARD, the balance of isoforms 1 and 3 might affect signaling pathways related to its kinase activity. For instance, hyperthermia may increase the alternative splicing kinetics or alter transcript stability and affect ERK pathways. A dominant-negative mechanism by the truncated isoform should not be excluded [41]. DNA-damaging or altered expression/subcellular distribution of RNA processing regulators [42] caused by temperature changes may also be responsible for the abnormal accumulation of alternatively spliced transcripts. These hypotheses obviously require experimental confirmation.

In the present study, *RIPK2* transcripts also showed a different expression pattern in samples from head and neck squamous cell carcinoma patients, with alternatively spliced transcript 2 exhibiting higher expression in tumors compared to their respective surgical margins. Differences in alternative splicing between tumor and normal samples have been described in the literature. For example, Gracio et al. [43], using ExonArray analysis of breast cancer and normal breast tissue samples, identified more than 200 genes with splicing differences associated with clinical outcome. Bjørklund et al. [44] obtained similar results by RNA-seq analysis in primary breast tumors for five genes. The large study of Kahles et al. analyzed 32 cancer types, including head and neck cancers, using RNA and whole-exome sequencing data and observed many differences in alternative splicing events in cancer compared with normal cells [45]. Specifically in regard to head and neck cancer, several other groups have described genes with differential expression of spliced variants [46-50]; among others). However, as far as we know, this is the first report showing differential expression of spliced transcripts of *RIPK2* gene between tumor and normal tissues.

At the protein level, decreased expression of the RIP-2 isoform 1 (immunoreactive band at ~61 kDa) was detected at low temperatures, disagreeing with our findings for full-length transcript 1, which showed no change at the same condition compared to the control. A divergent pattern was also observed for transcript 1 and the corresponding isoform 1 in neoplastic tissues, the former showing higher and the second lower levels in tumor samples compared with resection margins. This apparent discordance can be explained by the fact that protein abundance may differ from mRNA expression profile, mainly due to post-transcriptional control of gene expression or protein half-lives [51,52]. In addition, confirming our results from immunodetection assays, Wang and collaborators [12] also found reduced levels of RIP-2 in oral SCCs using immunohistochemical techniques.

In spite of using three distinct antibodies, another divergent result between RNA and protein expression was the absence of isoform 3, which suggests that translation of *RIPK2* into isoform 3 may not happen, at least in cell and tissue types analyzed in this study.

If the predicted alternative isoform 3 of RIP-2 is present in other conditions and tissue, then it should exhibit some impaired functions related to its kinase domain, including NFKB activation [5] [15,17,53], regulation of ERKs, p38 kinases, and own degradation [5,17,37,54].

Recently, Brady et al. identified a dominant-negative isoform of the translation initiation factor eIF-2B created by a hypoxia-mediated intron retention that inhibits translation and increases survival of neoplastic cells [50]. Dasgupta and collaborators [55] also described a C-terminal fragment of a member of RIP family, RIP-1, which can activate signaling events including NFKB and TNF pathways. They concluded that short RIP-1 with an aberrant N-terminal affects the long isoform levels and may represent a new regulation mechanism. Albeit exhibiting some differences, RIP-1 and −2 participate in the same regulatory pathway and therefore RIP-2 isoform also may have a similar function and modulate full-length RIP-2 under different conditions.

It is tempting to speculate that the Krieg ORF [18] could act as an uORF (upstream ORF) and repress translation of the downstream isoform 3 ORF, as cited for several stress response mRNAs by the literature [56,57]. The short intercistronic region between both ORFs should not be favorable for translation reinitiation due to an insufficient ribosomal scanning time necessary for reacquisition of the ternary complex (eIF2-GTP-Met-tRNAi) and for the downstream AUG recognition [58]. Translation reinitiation also depends on the Kozak context. Using numeric scores based on translation initiation site efficiencies determined in mammalian cells by Noderer et al.[59], both full-length and short RIP-2 transcripts have good efficiency values (90 and 107, respectively), but the predicted ratio of the initiation occurring at the second site compared to the first one is low (< 0.005), which may justify the absence of isoform 3.

As RIP kinases play a critical role in integrating stress signals, the elucidation of the factors that take part in the regulation of these proteins is of major importance. The alternative transcripts and protein isoforms may be tissue and context dependent and related to important disease responses [41].

Although the *RIPK2* transcript 2 has a coding potential, at present there is no direct evidence that it is translated in the isoform 3 or that it regulates biological processes, by competing with other molecules or modulating stress responses. Even without clarifying these issues, the present study raises many questions about RNA biology that may stimulate further functional investigation on the molecular mechanisms underlying *RIPK2* splicing regulation and their links to physiological and environmental changes.

## Conclusions

In conclusion, *RIPK2* transcripts 1 and 2 are expressed in different tissues and modulated by temperature, as determined by quantitative PCR assays. As far as we know, this is the first report showing splicing imbalances between tumor and normal samples of *RIPK2* gene, an important immune and inflammatory modulator. Despite transcription, no corresponding protein for the short isoform was detected in tissues and cell lines analyzed, which suggests that the balance of both transcripts may play regulatory roles and affect inflammation and other biological processes related to the activity of RIP-2.

## Acknowledgements

The authors thank the Fundação de Amparo à Pesquisa do Estãdo de São Paulo/FAPESP (FAPESP grant numbers 04/12054-9 and 10/51168-0) and Conselho Nacional de Pesquisas/CNPq (CNPq grant number 308904/2014-1) for financial support and fellowships. They are also grateful to Mauro Golin and Edilson Solim for artwork preparation, and to GENCAPO (Head and Neck Genome Project—http://www.gencapo.famerp.br/) team for the valuable discussions that motivated the present study.

